# TIR domains produce histidine-ADPR conjugates as immune signaling molecules in bacteria

**DOI:** 10.1101/2024.01.03.573942

**Authors:** Dziugas Sabonis, Carmel Avraham, Allen Lu, Ehud Herbst, Arunas Silanskas, Azita Leavitt, Erez Yirmiya, Mindaugas Zaremba, Gil Amitai, Philip J. Kranzusch, Rotem Sorek, Giedre Tamulaitiene

**Author notes:** These authors contributed equally (Dziugas Sabonis, Carmel Avraham).

## Abstract

TIR domains are central components of pattern recognition immune proteins across all domains of life. In both bacteria and plants, TIR-domain proteins were shown to recognize pathogen invasion and then produce immune signaling molecules exclusively comprising nucleotide moieties. Here we show that the TIR domain protein of the type II Thoeris defense system in bacteria produces a unique signaling molecule comprising the amino acid histidine conjugated to ADP-ribose (His-ADPR). His-ADPR is generated in response to phage infection and activates the cognate Thoeris effector by binding a Macro domain located at the C-terminus of the effector protein. By determining the crystal structure of a ligand-bound Macro domain, we describe the structural basis for His-ADPR recognition. Our analyses furthermore reveal a family of phage proteins that bind and sequester His-ADPR signaling molecules, allowing phages to evade TIR-mediated immunity. These data demonstrate diversity in bacterial TIR signaling and reveal a new class of TIR-derived immune signaling molecules combining nucleotide and amino acid moieties.

## Introduction

TIR (Toll/interleukin-1 receptor) domains are conserved protein domains essential for innate immunity in animals, plants, and bacteria^1,2^. These domains frequently form integral components of pattern recognition receptors, where their role is to initiate downstream immune signaling once infection is sensed^1,2^. It was recently shown that immune TIR domains in both plants and bacteria produce nicotinamide adenine dinucleotide (NAD^+^)-derived small signaling molecules that mediate the immune response, usually by inducing regulated death of the infected cell^2–5^. The extent of TIR signaling in plants and bacteria, and the repertoire of TIR-produced signaling molecules, is not completely understood.

Studies from the past few years revealed the role of TIR domain proteins in a bacterial anti-phage system called Thoeris^3,6–9^. In the Thoeris system of *Bacillus cereus* MSX-D12, TIR proteins first recognize phage infection, and then convert NAD^+^ into 1′′–3′ glyco-cyclic ADP ribose molecules (1′′–3′ gcADPR, also called 3′cADPR)^3,8,9^. 1′′–3′ gcADPR binds and activates a second, effector protein within the Thoeris system, called ThsA, which then depletes the cell of NAD^+^ and aborts the infection process^3,9^. It was further shown that some plant TIR-containing immune proteins synthesize gcADPR isomer molecules similar to those produced by bacterial Thoeris TIRs^3,4^.

Thoeris operons in bacterial genomes typically contain one or more TIR domain proteins (ThsB), each capable of recognizing a distinct set of phages, and a single *thsA* immune effector gene^3,6^. Two main architectures, called here type I and type II Thoeris, were described for bacterial Thoeris systems^3,6^ (Figure 1a). In type I systems typified by the well-studied Thoeris of *B. cereus*, ThsA contains a C-terminal SLOG domain that specifically binds 1′′–3′ gcADPR^7,8^ and an N-terminal sirtuin (SIR2) domain which is a potent NADase^3,7,10,11^. In type II Thoeris systems, the C-terminus of ThsA does not comprise a SLOG domain, and instead contains a Macro domain, a domain known to bind ADPR derivatives^12,13^ (Figure 1a). The N-terminus of ThsA in type II Thoeris comprises two transmembrane helices similar to other bacterial immune effectors that impair the bacterial cell membrane when activated^14^.

**Figure 1.**
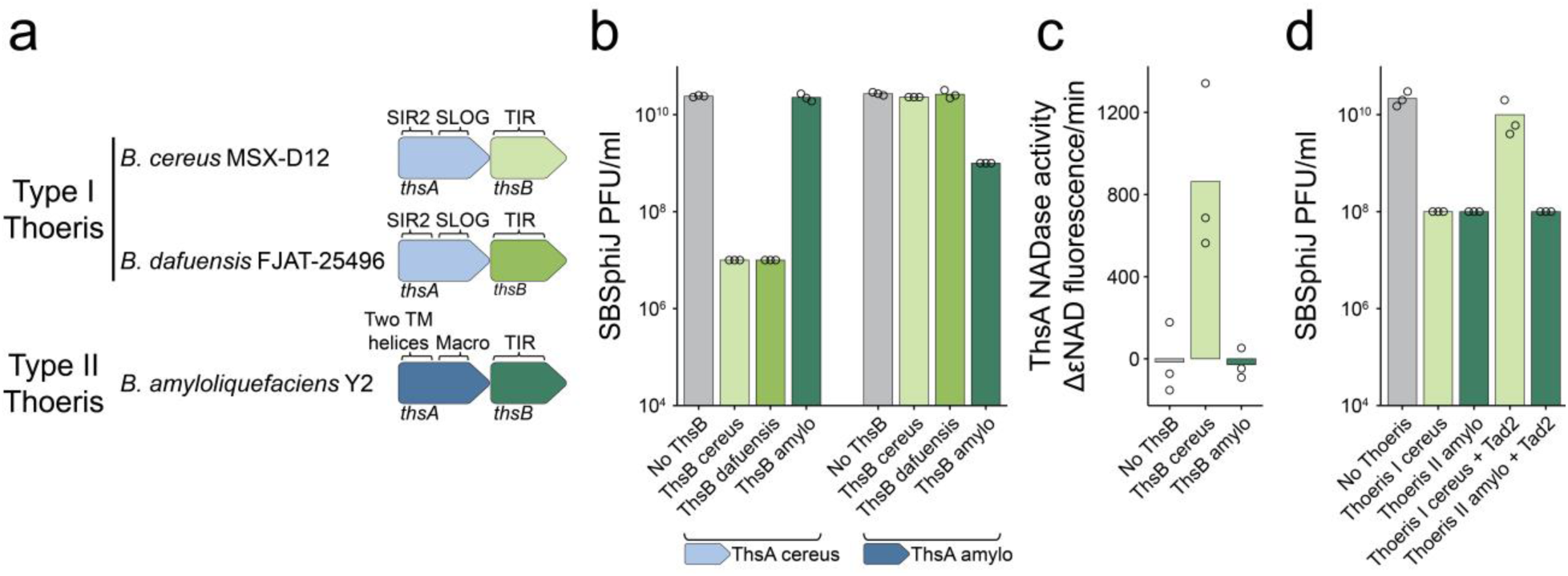
Two distinct types of Thoeris systems. **a**, Domain composition of the Thoeris systems studied here. **b**, Defense phenotypes in cells expressing combinations of ThsA and ThsB proteins. Data represent plaque-forming units per ml (PFU/ml) of SBSphiJ phage infecting cells that express the indicated ThsA and ThsB combinations. ThsA is expressed from its native promoter, while ThsB is under an IPTG-inducible promoter. “No ThsB” indicates control cells expressing GFP instead of ThsB. **c**, Activation of ThsA_cereus_ NADase activity by lysates from cells infected by phage SBSphiJ at multiplicity of infection (MOI) of 10. Infected cells express either the indicated ThsB or GFP as control (no ThsB). **d**, Tad2 inhibits type I but not type II Thoeris. Data represent PFU/ml of SBSphiJ phage infecting control cells (no Thoeris), cells expressing the type I Thoeris from *B. cereus* or type II Thoeris from *B. amyloliquefaciens*, and cells co-expressing a Thoeris system and the Tad2 protein from phage SPO1. Thoeris systems in this experiment are expressed from their native promoters. Bars in panels b, c, and d represent an average of three replicates, with individual data points overlaid.

In this study we characterized the type II Thoeris system from *Bacillus amyloliquefaciens* Y2^6^. Unexpectedly, we found that the TIR protein of type II Thoeris produces a new kind of signaling molecule comprising ADPR conjugated to the amino acid histidine. We show that the Macro domain of type II ThsA specifically binds the His-ADPR signaling molecule, and determine the structural basis for His-ADPR recognition by ThsA. Furthermore, we discover a family of phage proteins that specifically bind and sequester His-ADPR signals, enabling phages to overcome type II Thoeris defense.

## Results

### Type I and type II Thoeris systems produce different signaling molecules

The type I Thoeris systems from *B. cereus* MSX-D12 and from *Bacillus dafuensis* FJAT-25496 both protect against phage SBSphiJ^3^ (Figure 1a). Recombinant chimeric systems expressing the ThsA from *B. cereus* (ThsA_cereus_) and the ThsB TIR protein from *B. dafuensis* (TIR_dafuensis_) also defend against SBSphiJ, suggesting that TIRs from both these systems generate the same signaling molecule that activates the type I ThsA, as previously shown^3^ (Figure 1a,b).

The type II Thoeris system from *B. amyloliquefaciens* (alternatively called *B. velezensis*) also protects against phage SBSphiJ^6^ (Figure 1b, S1), suggesting that the TIR protein of type II also produces a signaling molecule in response to this phage. However, we found that chimeric systems expressing the ThsA protein from type II and the TIR protein from type I Thoeris, or vice versa, are incapable of defending against SBSphiJ (Figure 1b). These results imply that the signaling molecules produced by TIRs of one type of Thoeris cannot activate the ThsA of the second type.

To further determine whether TIRs from types I and II Thoeris systems produce similar or different molecules, we experimented with cells expressing the TIR domain protein alone, without the presence of the effector ThsA. We infected these cells with phage SBSphiJ, and then lysed the cells and filtered the lysates to enrich for small molecules. As expected, purified ThsA_cereus_ (from type I Thoeris) incubated with filtered cell lysates derived from infected TIR_cereus_- expressing cells exhibited strong NADase activity, confirming that the TIR_cereus_ protein from type I Thoeris produced 1′′–3′ gcADPR in response to SBSphiJ infection as previously demonstrated^15^ (Figure 1c). However, filtered cell lysates derived from cells expressing the TIR from type II Thoeris (TIR_amylo_) were not able to activate ThsA_cereus_ *in vitro*, confirming that the TIR protein of type II Thoeris does not produce a molecule capable of activating the ThsA from type I Thoeris.

A recent study reported that some phages encode an anti-Thoeris protein called Tad2 (Thoeris anti-defense 2), which binds and sequesters the 1′′–3′ gcADPR signaling molecule produced by Thoeris TIRs^15^. Tad2 forms a homotetrameric complex containing two pockets that bind gcADPR molecules with high affinity^15^. We co-expressed the type I Thoeris system with the Tad2 protein from phage SPO1, and found that Tad2 blocks Thoeris defense, as expected (Figure 1d). However, Tad2 did not abolish defense when co-expressed with type II Thoeris, further indicating that the type II system does not rely on the production of 1′′–3′ gcADPR molecules. As Tad2 is also known to bind and sequester the related molecule 1′′–2′ gcADPR^9^, our data suggest that the TIR protein of type II Thoeris generates neither 1′′–3′ gcADPR nor 1′′–2′ gcADPR.

### A phage protein sequesters the signaling molecule of type II Thoeris

Phylogenetic analyses of the Tad2 protein family revealed that homologs of this protein are encoded by phages infecting a large variety of bacteria spanning multiple taxonomic phyla^15^. Most of the Tad2 homologs that were previously tested experimentally were able to inhibit type I Thoeris, suggesting that they bind the 1′′–3′ gcADPR signaling molecule^15^ (Figure 2a). However, one Tad2 homolog, encoded by a prophage of *Myroides odoratus*, did not abolish antiphage defense when co-expressed with the type I Thoeris system^15^ (Figure 2a,b). We hypothesized that the Tad2 from *M. odoratus*, hereafter denoted ModTad2, evolved to recognize and sequester the signaling molecule of type II Thoeris. In support of this hypothesis, co-expression of ModTad2 with the type II Thoeris from *B. amyloliquefaciens* rendered this system unable to protect against phage SBSphiJ. Close homologs of ModTad2 from prophages of *Clostridium cadaveris* (CcaTad2) and *Tatumella morbirosei* (TmoTad2) were also capable of inhibiting type II Thoeris, suggesting that the clade of proteins represented by ModTad2 and its homologs can bind the molecule produced by the type II Thoeris TIR (Figure 2a,c, S2).

**Figure 2.**
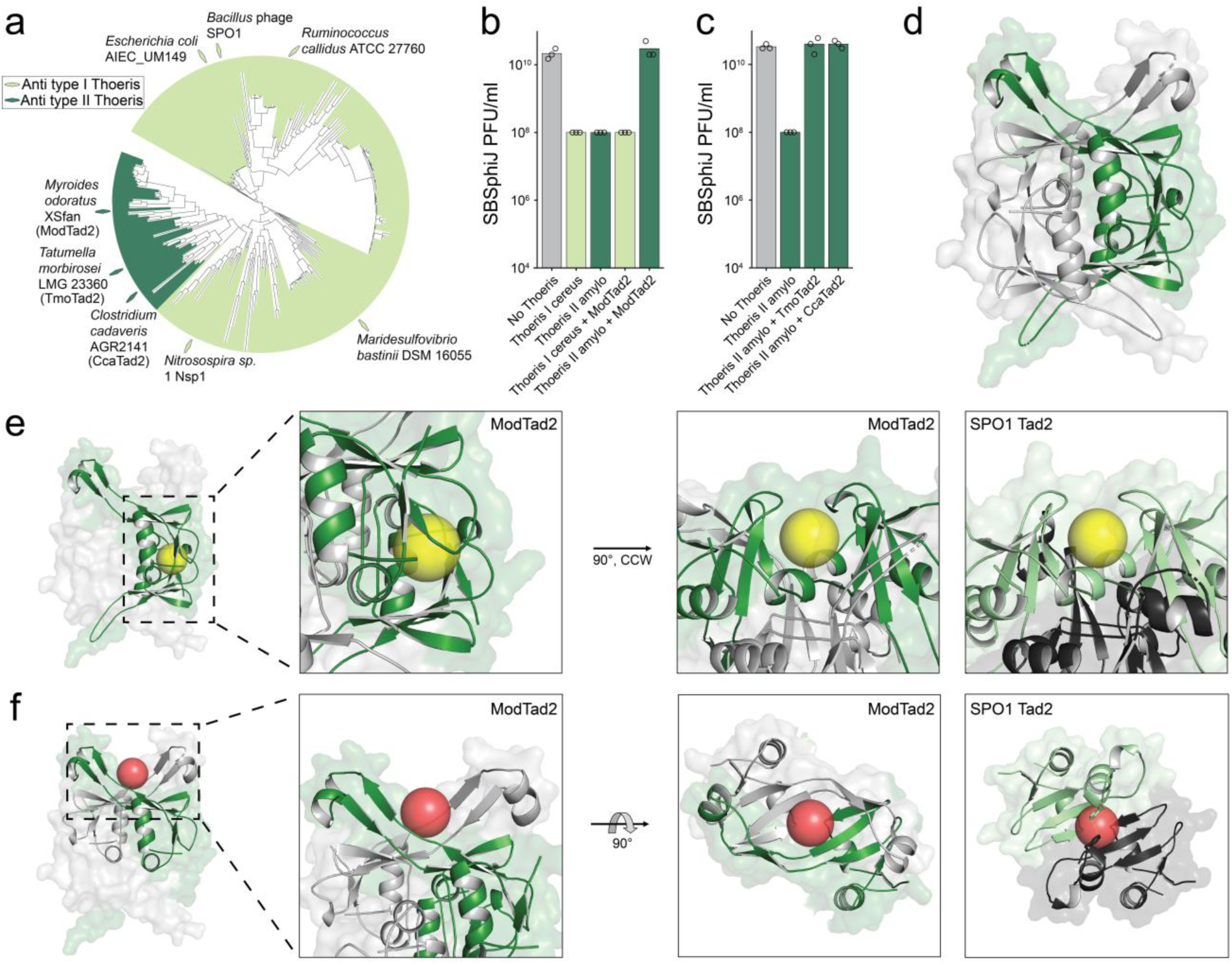
A phage protein that binds and sequesters the signaling molecule of type II Thoeris. **a**, Phylogenetic analysis of Tad2 homologs in phage and prophage genomes. The names of bacteria/phages from which Tad2 homologs were previously tested and found to inhibit type I Thoeris^15^ are indicated on the tree by light green diamonds. Names of Tad2 homologs found in the current study to inhibit type II Thoeris are indicated by dark green diamonds. Structure of presented tree is as in ref^15^. **b**, ModTad2 inhibits type II but not type I Thoeris. Data represent PFU/ml of SBSphiJ phage infecting bacterial cells as in Figure 1d. **c**, TmoTad2 and CcaTad2 inhibit type II Thoeris. **d**, Overview of ModTad2 structure. **e**, From left to right: Highlight of two ModTad2 monomers making up the canonical gcADPR binding site with a yellow sphere modeled in the cavity, zoom of the binding site, 90⁰ counter-clockwise rotation of the zoomed view, direct comparison of the same view for SPO1 Tad2, with one monomer in green and the other in grey. **f,** From left to right: Highlight of two ModTad2 monomers making up the putative novel binding site with a red sphere modeled centrally in the cavity, zoom of the putative binding site, 90⁰ inward rotation of the zoomed view, direct comparison of the same view for SPO1 Tad2 with one monomer in green and the other in grey. Ring stacking of F44 on each chain is highlighted in the SPO1 Tad2 view, showing possible occlusion of the pocket.

We determined the crystal structure of ModTad2 in the apo state. ModTad2 is a tetramer formed by two V-shaped homodimers, adopting an architecture similar to that of Tad2 from phage SPO1, which inhibits type I Thoeris defense^15^ (Figure 2d). While Tad2 from SPO1 exhibits two symmetric pockets that bind gcADPR, we observed four potential molecule-binding pockets in ModTad2 (Figure 2e,f). In addition to the two pockets that share structural homology with the gcADPR binding pocket of SPO1 Tad2, ModTad2 exhibits a potentially novel pocket within each V-shaped homodimer, formed by an extended region after strand β2, where an insertion of additional flanking beta strands results in extending helix α2 away from the core of the structure (Figure 2f). This pocket is occluded in SPO1 Tad2 where two phenylalanine residues (F44) stack, bringing the α2 helices together (Figure 2f). A recent study revealed that homologs of Tad2 can bind cyclic dinucleotide immune signaling molecules in the same position where we identified the new pocket in ModTad2^16^. It is possible that the signaling molecule of type II Thoeris is bound by either of these pockets.

To gain further insight into the nature of the signaling molecule produced by the type II Thoeris TIR in response to infection, we used ModTad2 as a “sponge” to bind this molecule. For this, we incubated ModTad2 with filtered lysates derived from phage-infected cells expressing TIR_amylo_, washed the ModTad2 complexes by successive dilution and concentration, and heated the ligand-bound ModTad2 at 98°C to denature the protein and release the signaling molecule (Figure 3a). Metabolomic analysis using untargeted mass spectrometry (MS) revealed a unique mass with a retention time of 9.04 min and an m/z value of 697.1374 (positive ionization mode) that was present in the sample retrieved from the denatured ModTad2 (Figure 3a). This mass was also present in filtered cell lysates prior to exposure to ModTad2, and was eliminated from these lysates following exposure to ModTad2, indicating that ModTad2 specifically binds and sequesters this molecule (Figure 3a). The unique molecule could not be detected in lysates derived from control cells expressing GFP instead of TIR_amylo_ (Figure 3a, S3), suggesting that this molecule is specifically produced by TIR_amylo_.

**Figure 3.**
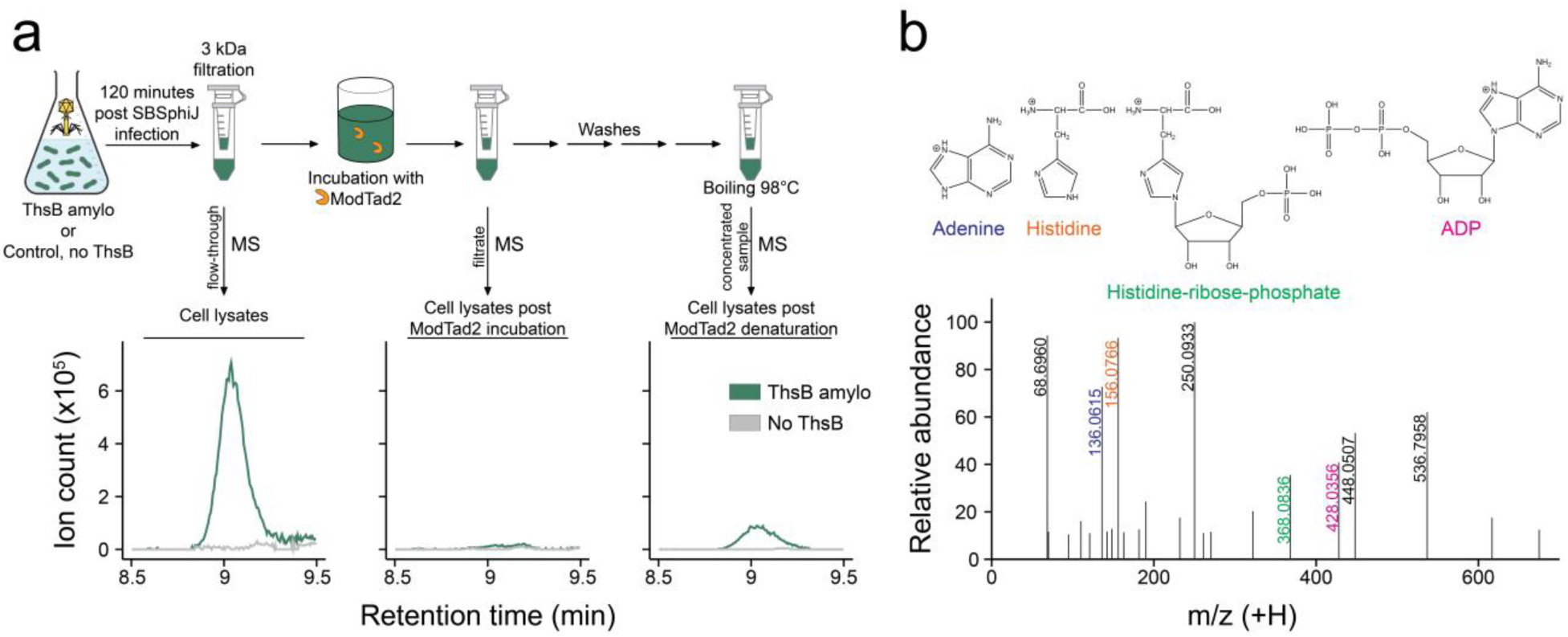
A small-molecule ligand bound by ModTad2. **a,**ModTad2 binds a small molecule ligand. Cells expressing ThsB_amylo_ or control cells that express GFP instead were infected with phage SBSphiJ at MOI of 10. After 120 min the cells were lysed, lysates were filtered, and then lysates were incubated with purified ModTad2. The lysates prior and post incubation with ModTad2, or after denaturation of ModTad2, were analyzed by mass spectrometry (MS). Extracted mass chromatograms of ions with an m/z value of 697.1374 and retention time of 9.04 min in positive mode are presented. Representative of two replicates, full replicate data are presented in Figure S3. **b**, MS/MS fragmentation spectra of the type II Thoeris-derived molecule. Hypothesized structures of selected MS/MS fragments are presented. The presented MS/MS data was obtained for the filtrate post ModTad2 denaturation in positive ionization mode.

Tandem mass spectrometry analysis (MS/MS) of the molecule released by ModTad2 revealed fragments with m/z values of adenine and adenine-di-phosphate (ADP), suggesting that the molecule may be related to ADPR (Figure 3b, Table S1). In addition to these fragments, however, we detected an abundant fragment with an m/z value of 156.0766, which, when compared to common cellular metabolites, unexpectedly revealed a perfect match to the amino acid histidine. Another abundant fragment ion exhibited an m/z value matching the expected mass of histidine- ribose-phosphate (Figure 3b). These data suggested that the signaling molecule involves derivatives of both adenine nucleotide and histidine (Figure 3b).

### Structure of ThsA Macro domain reveals a bound His-ADPR ligand

In the type II Thoeris system, the ThsA protein effector has a C-terminal Macro domain, which is predicted to bind the signaling molecule derived from the respective TIR protein (Figure 1a). To gain further insight into the new signaling molecule, we co-expressed TIR_amylo_ with the Macro domain of ThsA_amylo_ (residues 83–297, Macro_amylo_). We then determined a crystal structure of Macro_amylo_ bound to the ligand molecule at 2.23 Å resolution (Figure 4a). We found that the Macro domain of type II Thoeris forms a homodimeric complex, with each protomer possessing a globular α/β/α sandwich fold typical to Macro domain structures, composed of a central seven-stranded mixed β sheet (β1–β2–β12–β10–β3–β7–β6) flanked by seven α helices (Figure S4a). Dali structure comparison^17^ revealed that Macro_amylo_ exhibits similarity to an ADPribosyl hydrolase MacroD-like domain from the bacterium *Oceanobacillus iheyensis* and to a catalytically inactive MacroH2A-like domain from the ameboid protist *Capsaspora owczarzaki* (Figure S4b)^13,18,19^. Comparing Macro_amylo_ to these structures, we found that the loops near the ligand binding pocket are longer in Macro_amylo_ and possess an additional β hairpin (strands β4 and β5) and a small additional beta sheet (strands β8, 9, 11). In the ligand-binding pocket the adenine binding residues are conserved but the MacroD-type catalytic aspartate is absent in the Macro_amylo_ domain (Figure S4c). Although Macro domains are generally known to be monomeric^13,20^, Macro_amylo_ forms a dimer in the crystal with a significant dimer interface of ∼960 Å^2^ involving helices α4 and α5 (Figure 4b).

**Figure 4.**
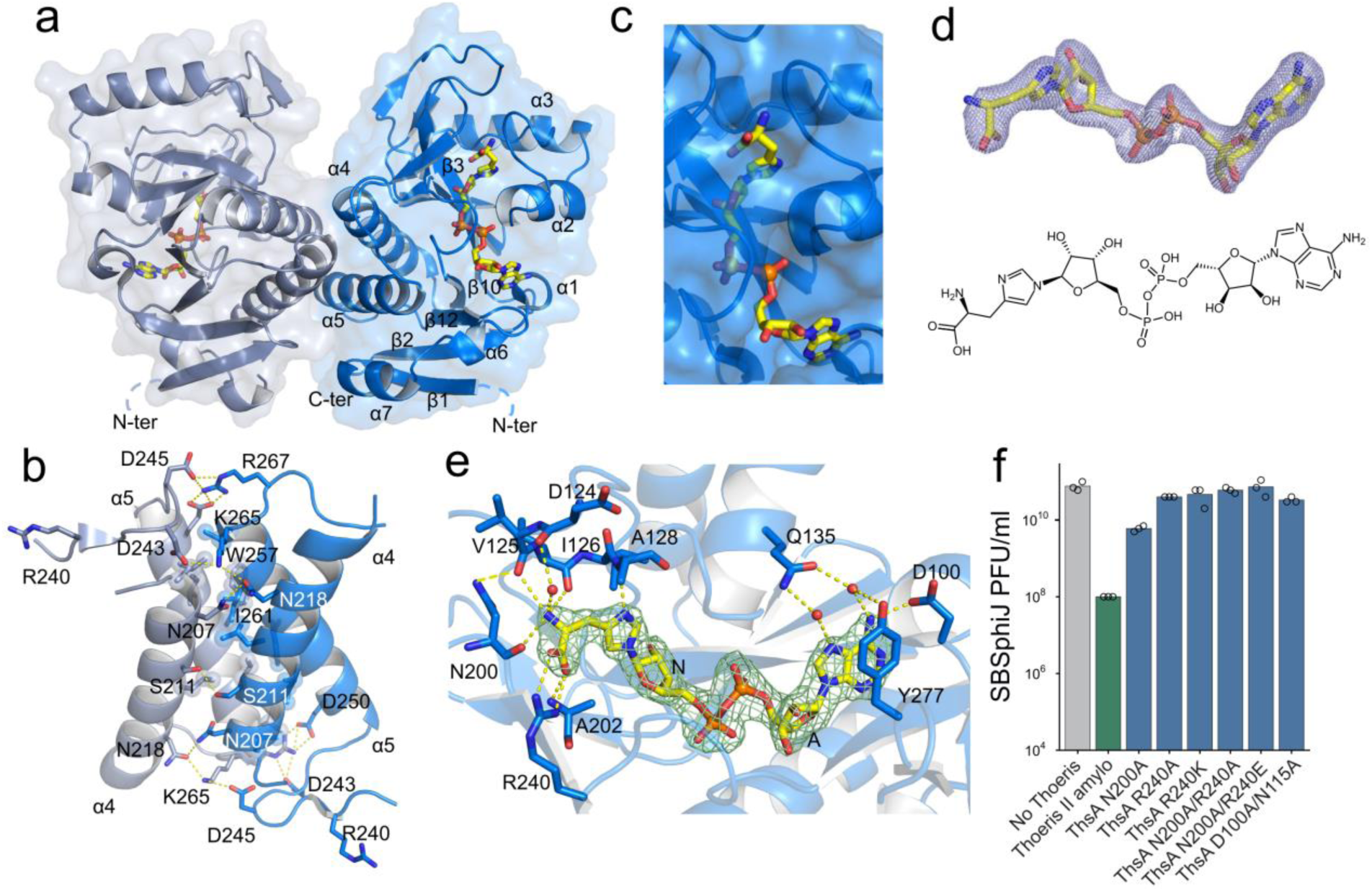
Structure of the ThsA_amylo_ Macro domain bound to His-ADPR. **a**, Macro_amylo_ domain dimer structure. Protomers are depicted in different colors, His-ADPR shown in yellow. **b**, Dimerization interface of the Macro_amylo_ domain. **c**, The His-ADPR binding pocket. **d**, Polder omit map of the ligand-binding site contoured at 5σ, showing the chemical structure of the ligand to be His-ADPR. **e**, Detailed view of the Macro_amylo_ residues interacting with His- ADPR adenine base and histidine moiety. Yellow dashed lines denote hydrogen bonding interactions. A and N marks A- and N-ribose, respectively. Light green mesh denotes His-ADPR 2F_o_ – F_c_ electron density contoured at 1.5σ. A full view of the Macro_amylo_ domain interactions with His-ADPR is presented in Figure S4d. **f**, Results of phage infection experiments of WT ThsA and the His-ADPR pocket mutants. Cells express either the WT Thoeris system, or a Thoeris system in which ThsA contains the indicated point mutation.

Further examining the predicted ligand-binding region in Macro_amylo_, we found that each ThsA Macro_amylo_ protomer has a ligand molecule bound in a deep elongated conserved binding pocket (Figure 4a,c). Clear electron density in the binding pocket, as well as a hydrogen bonding network with the protein, allowed us to identify the ligand bound in the Macro_amylo_ domain pocket as a histidine covalently linked to ADPR (termed here His-ADPR) (Figure 4d,e). In this molecule, the histidine is connected to the ADPR N-ribose‘s C1′ atom through its side chain NE1 (τ) atom (Figure 4d). The theoretical mass of His-ADPR in the protonated form (697.1379) is within the expected measurement error of the mass of the molecule we identified bound to ModTad2 (697.1374), and the MS/MS fragments determined for the ModTad2-derived molecule match the His-ADPR structure (Figure 3b). These results reveal that the type II Thoeris system utilizes His-ADPR as a signaling molecule.

Further analysis of the His-ADPR binding site demonstrated that the Macro_amylo_ domain makes numerous contacts with the His-ADPR ligand (Figure 4e, S4d). Binding of the adenine moiety is conserved with other Macro domains, involving stacking interaction with Y277 and a hydrogen bond between D100 and the N6 of the adenine base (Figure S4c,d)^13,18^. The Macro_amylo_ domain side chains Q135 and Y277 also make water-mediated hydrogen bonds to the N6 and N7 atoms of the adenine. The A-ribose and diphosphate of His-ADPR are bound mostly by protein backbone atoms, and the O2′ and O3′ of the N-ribose make contacts with the side chains of N133 and S194, respectively (Figure S4d). The histidine moiety of the His-ADPR ligand is bound by main chain atoms of Macro_amylo_, as well as by residue R240 of Macro_amylo_, which makes a side-chain contact with the histidine carboxylic oxygens. Mutations of the conserved Macro domain residues and the residues involved in His-ADPR binding abolished the antiphage activity of type II Thoeris (Figure 4f).

## Discussion

Our study establishes His-ADPR as the immune signaling molecule produced by the type II Thoeris defense system. This molecule represents a new class of TIR-derived immune signaling molecules comprising an amino acid conjugated to the nucleotide ADPR. Prior to the current study, immune TIR domain proteins in plants and bacteria were shown to produce exclusively nucleotide-based signaling molecules, in which ADPR was either cyclized^4,8,9^ or conjugated to another nucleotide^21^. In other cases, TIR domains were shown to produce nucleotide-based signals by processing the ends of DNA and RNA^22^. Our discovery raises the possibility that nucleotide-amino acid conjugates may serve as TIR-derived second messengers in other TIR-dependent defense systems.

Defense systems that rely on second messenger signaling are abundant in bacteria and were detected in at least 20% of sequenced bacterial and archaeal genomes^23^. In addition to Thoeris, these systems include CBASS^24–26^, Pycsar^27^ and type III CRISPR-Cas^28,29^. All these systems generate nucleotide-centric signaling molecules, with each system employing distinct enzymatic biochemistry for signal production: nucleotide polymerization in the case of CBASS and type III CRISPR systems, mononucleotide cyclization in Pycsar systems, and TIR-mediated NAD^+^ processing for Thoeris^30^. Interestingly, a recent study discovered a version of type III CRISPR-Cas that utilizes the nucleotide polymerization biochemistry of Cas10 to conjugate S-adenosyl methionine (SAM) to ATP, generating a SAM-AMP immune signaling molecule^31^. Together with the results of our study, these data demonstrate that enzymes producing immune signaling molecules can evolve to incorporate non-nucleotide moieties within the produced signal. As phages express nuclease and sponge proteins that degrade and sequester specific immune signaling molecules^9,15,32–35^, this generates a strong evolutionary pressure for signal diversification in bacterial defense systems.

For many years, TIR domains were known only as protein-protein interaction modules^36^, and the NADase activity of TIR domains was discovered relatively recently^37–39^. TIR-mediated small molecule signaling in immunity is a very recent discovery, with the identity of the first TIR-derived molecules determined only in 2022^8,9^. So far, TIRs in plants and bacteria were shown to produce diverse molecules including gcADPR^8,9^, pRib-AMP/ADP^40^, ADPR-ADPR^21^, ADPR-ATP^21^, and now His-ADPR. Combined together, these discoveries demonstrate remarkable plasticity in the ability of TIR domains to produce signaling molecules via conjugation of ADPR and suggest that more TIR-produced signaling molecules may be discovered in the future.

## Methods

### Bacteria growth conditions

For the experiments presented in Figures 1, 2, S1 and S3, bacteria were grown in MMB (LB supplemented with 0.1 mM MnCl_2_ and 5 mM MgCl_2_) in liquid medium with shaking at 200 rpm at 37°C or 25°C as stated in Table S2, or on LB 1.5% agar plates. The antibiotics spectinomycin (100 µg/ml) or chloramphenicol (5 µg/ml) were used to ensure the presence of an integrated antibiotics resistance cassette in the *B. subtilis* BEST7003 genomic *amyE* or *thrC* locus respectively. When applicable, 1 mM or 0.1 mM isopropyl β-d-1-thiogalactopyranoside (IPTG) was added to bacterial cultures to induce gene expression as stated in Table S2. A list of all bacterial strains and phages used in this study can be found in Table S2 and a list of all plasmids and primers used in this study can be found in Table S3.

### Cloning and transformation

The shuttle vectors used in this study, as well as the DNA for the Thoeris genes or Thoeris anti-defense genes were constructed in previous studies^6,9^. Thoeris defense systems or Thoeris ThsA genes were cloned under native promoters in the shuttle vector pSG1-rfp^6^ that contains the spectinomycin-resistance gene, and the cloned sequence with the spectinomycin-resistance gene was integrated into the *B. subtilis amyE* locus. As a negative control, a transformant with an empty insert containing only the spectinomycin-resistance gene in the *amyE* locus, was used.

Anti-Thoeris genes or Thoeris ThsB genes were cloned under an IPTG inducible promoter (Phspank) in the shuttle vector pSG-thrC-phSpank^9^ that contains the chloramphenicol-resistance gene, and the cloned vector was integrated into *B. subtilis thrC* locus in the appropriate background (Table S2). As a negative control, a transformant with an identical plasmid, containing GFP instead of an anti-Thoeris gene or a *thsB* gene, was used and integrated in the *thrC* locus.

To generate the plasmids used in this study, genes were amplified using KAPA HiFi HotStart ReadyMix (Roche cat # KK2601), cloned using NEBuilder HiFi DNA Assembly cloning kit (NEB, cat # E5520S) and transformed to NEB 5-alpha Competent *E. coli* High Efficiency (NEB). For one-fragment DNA assembly, the linear plasmid obtained by PCR was ligated using the KLD enzyme mix (NEB cat # M0554S) according to the manufacturer’s protocol before transformation. For assembly of more than one fragment, PCR products were treated with the FastDigest DpnI (ThermoFisher) restriction enzyme according to the manufacturer’s protocol. The fragments were then assembled using the NEBuilder HiFi DNA Assembly Master Mix before transformation Plasmids purified from an overnight culture were then transformed into *B. subtilis* BEST7003 cells. Transformation was performed using MC medium as previously described^6^.

### SBSphiJ propagation

Overnight liquid cultures of *B. subtilis* cells were diluted 1:100 in 100 ml MMB, and grown at 25°C, 200 rpm shaking to an OD_600_ of 0.3. At this stage, Phage SBSphiJ was added to the liquid culture at an MOI of 0.1 and incubation at 25°C, 200 rpm shaking continued until culture collapse. The culture was then centrifuged at 4°C for 10 min at 3200 *g* and the supernatant was filtered through a 0.22 µm filter to get rid of remaining bacteria and large debris.

### Plaque assays

Phage SBSphiJ titer was determined using the small drop plaque assay method^41^. 400 μl of an overnight culture of bacteria grown in MMB with antibiotics were mixed with 30 ml pre-melted 0.5% MMB agar and poured into a 10 cm square plate. For induction of genes expressed under the Phspank promoter, IPTG was added to a final concentration of 1 mM or 0.1 mM before plating (see Table S2). After incubation for 1 h at room temperature, 10 µl drops from 10-fold serial dilutions of the phage lysate in MMB were dropped on top of the bacterial layer. After the drops dried up, plates were incubated at 25°C overnight. Plaque forming units (PFU) were determined by counting the derived plaques, and lysate titer was determined by calculating PFU/ml. When no individual plaques could be identified, a faint lysis zone across the drop area was considered to be 10 plaques. Bacterial defense phenotype was measured as the ratio between the PFU/ml on control bacteria and PFU/ml on bacteria expressing a Thoeris system.

### Phage-infection dynamics in liquid medium

Overnight cultures of *B. subtilis* cells expressing a Thoeris system or an empty insertion were diluted 1:100 in MMB medium supplemented with spectinomycin (100 μg/ml). Cultures were incubated at 25°C with shaking 200 rpm until cells reached OD_600_ of 0.3. At this point, 180 μl of the culture was transferred into a 96-well plate containing additional 20 μl of MMB for the uninfected control or phage SBSphiJ for a final MOI of 0.1 or 10. Plates were incubated at 25°C with shaking in a TECAN Infinite200 plate reader and absorbance at OD_600_ was measured every 4 minutes.

### NADase activity assay with purified ThsA_cereus_

Lysates for this assay were prepared from an overnight culture of *B. subtilis* cells expressing ThsB_cereus_, ThsB_amylo_ or control cells expressing GFP. Cells were diluted 1:100 in 100 ml MMB medium supplemented with 1 mM IPTG and chloramphenicol (5 µg/ml), and grown at 25°C, 200 rpm shaking until reaching an OD_600_ of 0.3. At this point, SBSphiJ phage was added at an MOI of 10 and cultures were incubated at 25°C for 120 min. Then, 50 ml samples were collected and centrifuged at 4°C, 3200 *g* for 10 min to pellet the cells. The supernatant was discarded, and the pellet was flash frozen and stored at −80°C.

To extract the cell metabolites from frozen pellets, 600 μl of 0.1M Na phosphate buffer, pH 8.0, was added to each pellet and incubated at room temperature for 10 min, and then transferred to ice. Then, the samples were transferred to a FastPrep Lysing Matrix B in a 2 ml tube (MP Biomedicals cat # 116911100) and lysed at 4°C using a FastPrep bead beater for 2 × 40 s at 6 m/s. Tubes were then centrifuged at 4°C for 10 min at 15,000 *g*. Supernatant was transferred to Amicon Ultra-0.5 Centrifugal Filter Unit 3 kDa (Merck Millipore cat # UFC500396) and centrifuged for 45 min at 4°C, 12,000 *g*. Filtered cell lysates were taken for in vitro ThsA_cereus_ activity assay.

ThsA_cereus_ was expressed and purified as described in a previous study^15^. The NADase reaction was performed in black 96-well half area plates (Corning, cat #3694). In each reaction well, purified 2 µl ThsA_cereus_ protein was added to 43 µl filtered cell lysates (final concentration 100 nM). 5 µl of 5 mM nicotinamide 1,N^6^-ethenoadenine dinucleotide (εNAD^+^, Sigma cat# N2630) was added to each well immediately before measurements. Plates were incubated inside a Tecan Infinite M200 plate reader at 25°C, and measurements were taken at 300 nm excitation wavelength and 410 nm emission wavelength every 15 s. Reaction rate was calculated from the linear part of the initial reaction.

### Preparation of filtered cell lysates for LC-MS analysis

For generating filtered cell lysates that contain TIR-catalyzed signaling molecules, we used *B. subtilis* cells expressing ThsB_amylo_ or GFP for control, under an inducible Phpsank promoter. These cultures were grown overnight and then diluted 1:100 in 250 ml MMB medium supplemented with 1 mM IPTG and chloramphenicol (5 µg/ml) and grown at 37°C, 200 rpm shaking for 90 min, followed by additional incubation and shaking at 25°C, 200 rpm until reaching an OD_600_ of 0.3. At this point, SBSphiJ phages were added at an MOI of 10, and cultures were incubated at 25°C for 120 min. Then, 200 ml samples were collected and centrifuged at 4°C, 3200 *g* for 10 min to pellet the cells. The supernatant was discarded, and the pellet was flash frozen and stored at −80°C. Cell metabolites were extracted as mentioned above for ThsA_cereus_ activity assay, and filtered cell lysates were sent to LC-MS analysis.

### Isolation of type II Thoeris amylo signaling molecule for LC-MS analysis

To isolate the ThsB_amylo_ signaling molecule, 100 µl filtered cell lysates derived from cells expressing ThsB_amylo_ or GFP were incubated at room temperature for 1 hr with 75 µl 0.1 M Na phosphate buffer, pH 8 and 25 µl purified ModTad2 protein (67 µM, 0.66 mg/ml) to capture the molecule. Following incubation, the mixture was transferred to Amicon Ultra-0.5 Centrifugal Filter Unit 3 kDa and centrifuged for 20 min at 4°C, 12,000 *g*. This filtrate was sent to LC-MS analysis. To remove the metabolites that did not bind ModTad2, the columns were washed 4 times by adding 400 µl 0.1 M Na phosphate buffer, pH 8, and centrifuged for 20 min at 4°C, 12,000 *g*. ModTad2 was recovered by flipping the 3 kDa filter, transferring it to a new tube, and centrifuging it for 5 min at 4°C, 1,000 *g*. To collect the ModTad2-bound molecules, ModTAD2 was denatured for 25 min at 98°C. To remove the ModTad2 protein, the sample was filtered in an Amicon Ultra-0.5 Centrifugal Filter Unit 10 kDa (Merck Millipore cat # UFC501096) supplemented with a 100 µl of 0.1 M Na phosphate buffer, pH 8, and centrifuged 20 min at 4°C, 12,000 *g*. The purified molecules were sent to LC-MS analysis.

### LC-MS analysis

Prior to the LC-MS analysis, samples were centrifuged twice at 18,000 *g* to remove possible precipitants and transferred to a high-performance liquid chromatography (HPLC) vial. Samples were analyzed as described previously^42^ with minor modifications described below. Briefly, analysis was performed using Acquity I class UPLC System combined with mass spectrometer Q Exactive Plus Orbitrap™ (Thermo Fisher Scientific), which was operated in positive and negative ionization modes. The MS spectra were acquired with 70.000 resolution, scan range of 400 – 2000 m/z. For the identification of the compounds, we used a data-dependent acquisition, top 5 method. The LC separation was done using the SeQuant ZIC-pHILIC (150 mm × 2.1 mm) with the SeQuant guard column (20 mm × 2.1 mm) (Merck). The mobile phase B was acetonitrile and the mobile phase A was 20 mM ammonium carbonate with 0.1% ammonia hydroxide in a 80:20 solution (v/v) of double-distilled water and acetonitrile. The flow rate was kept at 200 μl/min, and the gradient was as follows: 75% of B (0-2 min), decreased to 25% of B (2-14 min), 25% of B (14-18 min), increased to 75% of B (18-19 min), 75% of B (19-23 min). The data was analyzed using MZmine 2.5.3 software.

Untargeted mass spectrometry (MS) data from all samples was integrated, and the signaling molecule generated by TIR_amylo_ was identified by searching for molecules enriched in TIR_amylo_ filtered cells lysates prior to ModTad2 incubation and post ModTad2 denaturation but absent in TIR_amylo_ filtered cell lysates post ModTad2 incubation and control cells that lack TIR_amylo_. The analysis was performed for the MS measurements derived from both positive and negative ionization modes. To define the m/z value and retention time of His-ADPR, the same analysis as before was repeated on the MS results of TIR_amylo_ filtered cells lysates, with the exception that the parameters in MZmine “Chromatogram deconvolution” feature were adjusted to engulf the same peak in two repeats.

### ModTad2 protein expression and purification for biochemistry

Expression of ModTad2 was performed using the expression vector pET28-bdSumo. This vector was constructed by transferring the His14-bdSUMO cassette from the expression vector (Designated K151) generously obtained from Prof. Dirk Görlich from the Max-Planck-Institute, Göttingen, Germany^43^ into the expression vector pET28-TevH^44^. Cloning was performed by the Restriction-Free (RF) method^45^. A 5 l culture of BL21(DE3) harboring the vector was induced with 200 µM IPTG and grown at 15°C overnight. The cells were harvested and lysed by a cell disrupter (Constant Systems) in buffer comprising 50 mM Tris pH 8, 0.5 M NaCl, 30 mM imidazole, 1 mM MgCl_2_, and containing 200 KU/100 ml lysozyme, 20 μg/ml DNase, 1 mM phenylmethylsulfonyl fluoride (PMSF) and protease inhibitor cocktail. After clarification of the supernatant by centrifugation, the lysate was incubated with 5 ml pre-washed Ni^2+^ beads (Adar Biotech) for 1 h at 4°C. After removing the supernatant, the beads were washed 4 times with PBS buffer. The cleaved protein (without tags) eluted from the beads by incubation of the beads with 5 ml cleavage buffer: PBS supplemented with 250 mM sucrose and 10% glycerol containing 0.1 mg bdSumo protease (without His tag) for 2 h at RT. The supernatant containing the cleaved protein was removed and applied to a size exclusion (SEC) column (HiLoad_16/60_Superdex75 prep- grade, GE Healthcare) equilibrated with PBS. The pure protein was pooled and frozen in aliquots stored at -80°C.

### Expression and purification of ModTad2 for crystallization

ModTad2 was codon optimized (GeneArt), synthesized as DNA fragments (Integrated DNA Technologies), and cloned by Gibson assembly into a custom pET vector with an N-terminal 6× His-SUMO tag and an ampicillin resistance gene. The plasmid was transformed into BL21(DE3) RIL *E. coli* (Agilent), colonies were grown on 1.5% agar MDG plates (2 mM MgSO_4_, 0.5% glucose, 25 mM Na_2_HPO_4_, 25 mM KH_2_PO_4_, 50 mM NH_4_Cl, 5 mM Na_2_SO_4_, 0.25% aspartic acid and 2–50 μM trace metals, 100 µg/ml ampicillin and 34 µg/ml chloramphenicol), and 3 colonies were picked to inoculate separate 30 ml MDG starter cultures, which were grown overnight at 37°C with 230 rpm shaking. 1 l of M9ZB expression culture (2 mM MgSO_4_, 0.5% glycerol, 47.8 mM Na_2_HPO_4_, 22 mM KH_2_PO_4_, 18.7 mM NH_4_Cl, 85.6 mM NaCl, 1% Cas-amino acids, 2–50 μM trace metals, 100 µg/ml ampicillin and 34 µg/ml chloramphenicol) was seeded with 15 ml MDG starter culture, grown to OD_600_ of 2.5 at 37°C with 230 rpm shaking, and induced with 0.5 mM IPTG and lowering of temperature to 16°C. After 16-20 h, 2 l of culture was harvested by centrifugation, resuspended in 120 ml of lysis buffer (20 mM HEPES-KOH pH 7.5, 400 mM NaCl, 30 mM imidazole, 10% glycerol and 1 mM DTT), lysed by sonication, and clarified by centrifugation at 25,000 *g* for 20 min. Supernatant was passed over 8 ml of Ni-NTA resin (Qiagen), and the resin was washed with 70 ml of lysis buffer supplemented to 1 M NaCl followed by 20 ml lysis buffer. Protein was eluted with 20 ml of lysis buffer supplemented to 300 mM imidazole and dialyzed overnight at 4°C using 14 kDa dialysis tubing in size-exclusion buffer (20 mM HEPES-KOH pH 7.5, 250 mM KCl and 1 mM TCEP) in the presence of recombinant human-SENP2 to induce SUMO-tag cleavage. Protein was further purified by size-exclusion chromatography on a Superdex 75 16/600 column (Cytiva). Peak fractions were collected, concentrated to >50 mg/ml, flash frozen in liquid nitrogen, and stored at −80°C.

### ModTad2 crystallography and structural analysis

Crystals of ModTad2 were grown by the hanging drop vapor diffusion method using EasyXtal 15 well trays (NeXtal). First, protein was prepared at 10 mg/ml in crystallization buffer (20 mM HEPES-KOH pH 7.5, 80 mM KCl and 1 mM TCEP). 2 μl hanging drops of 1 μl protein and 1 μl reservoir solution were set above 400 μl of reservoir solution (0.1 M MgCl_2_, 0.1 M sodium acetate pH 4.6, 25% w/v PEG 400). Crystals were grown for 1 week before harvesting by flash freezing in liquid nitrogen. X-ray diffraction data were collected at the Advanced Photon Source (beamlines 24-ID-C and 24-ID-E), and data were processed using the SSRL autoxds script (A. Gonzalez, Stanford SSRL). Phases were determined by molecular replacement using a truncated predicted structure of ModTad2 (apo) from ColabFold v1.5.3^46^. Model building was performed in WinCoot^47^, with refinement in Phenix, and statistics were analyzed as presented in supplementary Table S4^48–50^). Final structures were refined to stereochemistry statistics for Ramachandran plot (favoured/allowed), rotamer outliers, and MolProbity score as follows: ModTad2 apo, 98.7%/1.3%, 3.4%, 1.74. See supplementary Table S4 and Data Availability for deposited PDB codes. All structure figures were generated with PyMOL 2.5.0.

### Expression and purification of ThsA_amylo_ Macro domain

Expression of ThsA_amylo_ Macro domain (Macro_amylo_, residues 83-297, with C-terminal TwinStrep- tag) was performed using the expression vector pBAD_DelTM-ThsA-TwinStrep_ThsB-His. To generate the pBAD_DelTM-ThsA-TwinStrep_ThsB-His expression vector, the type II Thoeris amylo operon was modified by removing the TM domain of ThsA_amylo_ [residues 1-82] and genetically fusing a C-terminal Twin-Strep tag. In addition, a C-terminal His_6_ tag was fused to the C-terminus of ThsB. The coding sequence of the operon was codon optimized for *E.coli* and synthesized by TWIST Biosciences. The operon was then cloned into the NcoI/HindIII site of a pBAD-His backbone by TWIST Biosciences. A 5 l culture of *E. coli* TOP10 cells harboring the vector was induced with 0.2% L-arabinose and grown overnight at 16°C. The cells were harvested by centrifugation and resuspended in the Purification buffer (20 mM Tris-HCl (pH 8.0 at 25°C), 1 M NaCl, 5 mM 2-mercaptoethanol, 0.1% TRITON X-100) supplemented with 2 mM phenylmethylsulfonyl fluoride (PMSF) and 5% (v/v) glycerol and lysed by sonication. After removing cell debris by centrifugation, the supernatant was loaded on Strep-Tactin XT Superflow column (IBA) and the bound protein was eluted with 50 mM D-biotin solution in the Purification buffer. Fractions with the protein of interest were pooled, concentrated up to 5 ml using Amicon Ultra-15 centrifugal filter unit (Merck Millipore) and loaded on a HiLoad 16/600 Superdex 200 gel filtration column (Cytiva) equilibrated with the Purification buffer containing 0.01% TRITON X-100. Peak fractions containing the protein of interest were pooled. Purified protein was dialyzed against 20 mM Tris-HCl (pH 8.0 at 25°C), 500 mM NaCl, 2 mM DTT, 0.01% TRITON X-100 and 50% v/v glycerol containing buffer and stored at -20°C. The final protein concentrations were determined by measuring absorbance at 280 nm using sequence-predicted extinction coefficients.

### Crystallization and structure determination of the Macro_amylo_ domain

Macro_amylo_ domain initial needle-shaped crystals were obtained by the sitting-drop vapor diffusion method from 6.9 mg/ml protein solution in the Concentration buffer (20 mM Tris-HCl (pH 8.0 at 25°C), 250 mM NaCl, 2 mM DTT, 0.01 % TritonX-100) mixed in 7:3 ratio with reservoir solution (25% (w/v) polypropylene glycol (PEG) 10000, 0.1 M Tris-acetate (pH 8.0 at 25°C), 0.1 M KCl, 0.05 M magnesium formate) at 20°C. The needle-shaped crystals were used for microseeding. Crystals, used for a structure determination, were obtained by the sitting-drop vapor diffusion method from 5 mg/ml protein solution in the Concentration buffer, reservoir solution (23.75% PEG 3500, 0.1 M Bis-Tris (pH 6.5 at 25°C), 0.19 M ammonium sulfate, 5% (v/v) glycerol) and seeding solution mixed in a 4:3:1 ratio.

The X-ray diffraction dataset was collected to the nominal resolution of 2.23 Å at the EMBL/DESY Petra III P13 beamline (Hamburg, Germany) at 100 K, wavelength 0.9800 Å. XDS^51^, SCALA and TRUNCATE^52^ were used for data processing. The structure was solved by molecular replacement by Phaser^53^ using an ensemble of 5 models obtained from Robetta server^54^ and rebuilt by Phenix AutoBuild^55^. The model was improved by several cycles of refinement in Phenix (phenix-1.20.1- 4487)^56^ and manual inspection in Coot 0.9.7^47^. Cif file for His-ADPR refinement was prepared using eLBOW^57^, bond lengths were corrected using data from^58^ and His.cif from Coot. C-terminal residues of the domain are disordered; final model contains residues 83-293 in chain A and 83- 289 in chain B. 96.38 % of the residues are in the favored and 3.62 % in the allowed region of the Ramachandran plot. The data collection and refinement statistics are presented in Table S4. The molecular graphics figures were prepared with PyMOL (v.2.3.0) (The PyMOL Molecular Graphics System, Schrödinger, LLC).

## Data Availability

All data are available in the Article and the Supplementary Material. Strains of bacteria and phages and plasmid maps of the constructs used for the experiments are attached as Supplementary Files. The atomic coordinates and structure factors have been deposited in the Protein Data Bank under accession codes 8V3E (ModTad2) and 8R66 (His-ADPR bound Macro_amylo_).

## Acknowledgements

We thank Orly Izsak for help with mass spectrometry analysis, and Maxim Itkin and Sergey Malitsky from the metabolic profiling unit of the Weizmann Institute of Science for conducting the MS experiments. We thank Yoav Peleg and Shira Albeck from the Center for Structural Proteomics of the Weizmann Institute of Science for assistance with ModTad2 purification. We thank the Sorek laboratory members for comments on earlier versions of this manuscript. X-ray diffraction data were collected at the beamline P13 operated by EMBL Hamburg at the PETRA III storage ring (DESY, Hamburg, Germany) and at the Northeastern Collaborative Access Team beamlines 24-ID-C and 24-ID-E (P30 GM124165). Access to the EMBL P13 beamline has been supported by iNEXT-Discovery, project number 871037, funded by the Horizon 2020 program of the European Commission. NE-CAT data collection was supported by use of a Pilatus detector (S10RR029205), an Eiger detector (S10OD021527), and the Argonne National Laboratory Advanced Photon Source (DE-AC02-06CH11357). R.S. was supported, in part, by the European Research Council (grant no. ERC-AdG GA 101018520), Israel Science Foundation (MAPATS Grant 2720/22), the Deutsche Forschungsgemeinschaft (SPP 2330, Grant 464312965), the Ernest and Bonnie Beutler Research Program of Excellence in Genomic Medicine, Dr. Barry Sherman Institute for Medicinal Chemistry, Miel de Botton, the Andre Deloro Prize, and the Knell Family Center for Microbiology. P.J.K. was supported, in part, by the Pew Biomedical Scholars program and The Mathers Foundation. G.T. was supported by the Research Council of Lithuania (LMTLT) (grant S-MIP-21-6). E.H. was supported by the Israel Cancer Research Fund Postdoctoral Fellowship. E.Y. was supported by the Clore Scholars Program, and, in part, by the Israeli Council for Higher Education (CHE) via the Weizmann Data Science Research Center.

---

**Figure S1.**
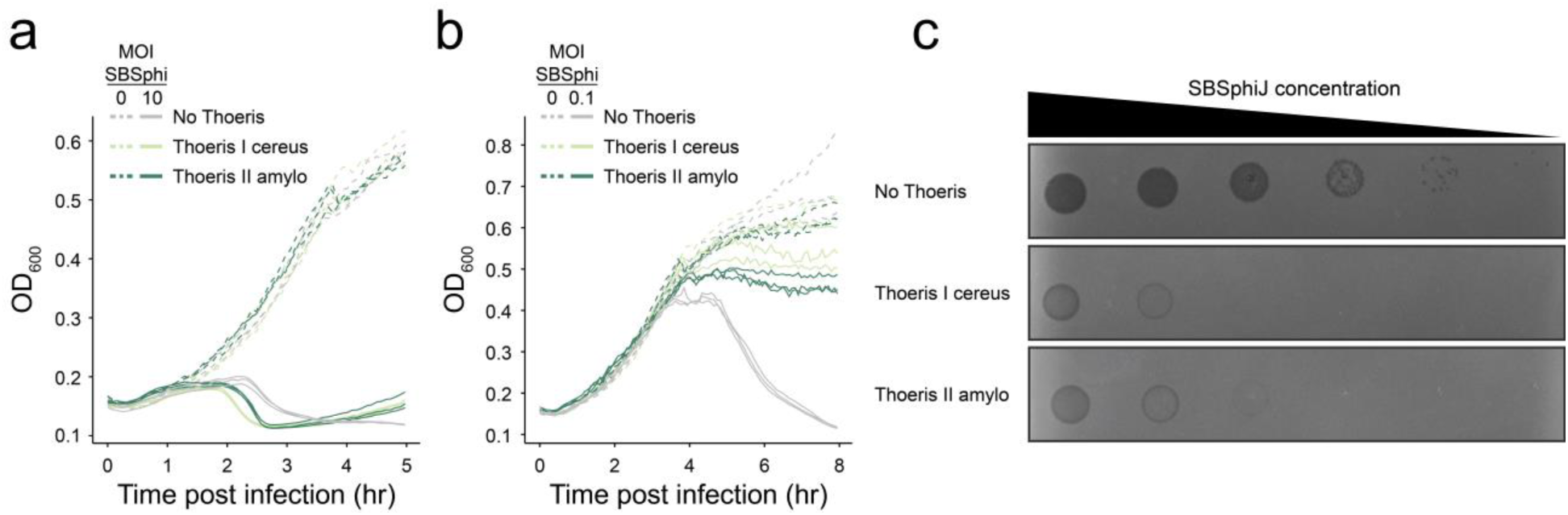
Both type I and type II Thoeris systems protect via abortive infection. **a**, Growth curves of Thoeris- expressing and control cultures with and without infection by phage SBSphiJ at an MOI of 10. Data from three replicates are presented as individual curves. OD_600_, optical density at 600 nm. **b**, Same as in panel A, but with cells infected at MOI of 0.1. **c**, A representative plaque assay showing that both type I and type II Thoeris systems protect from phage SBSphiJ. Shown are tenfold serial dilution plaque assays with phage SBSphiJ. Representative of three replicates.

**Figure S2.**
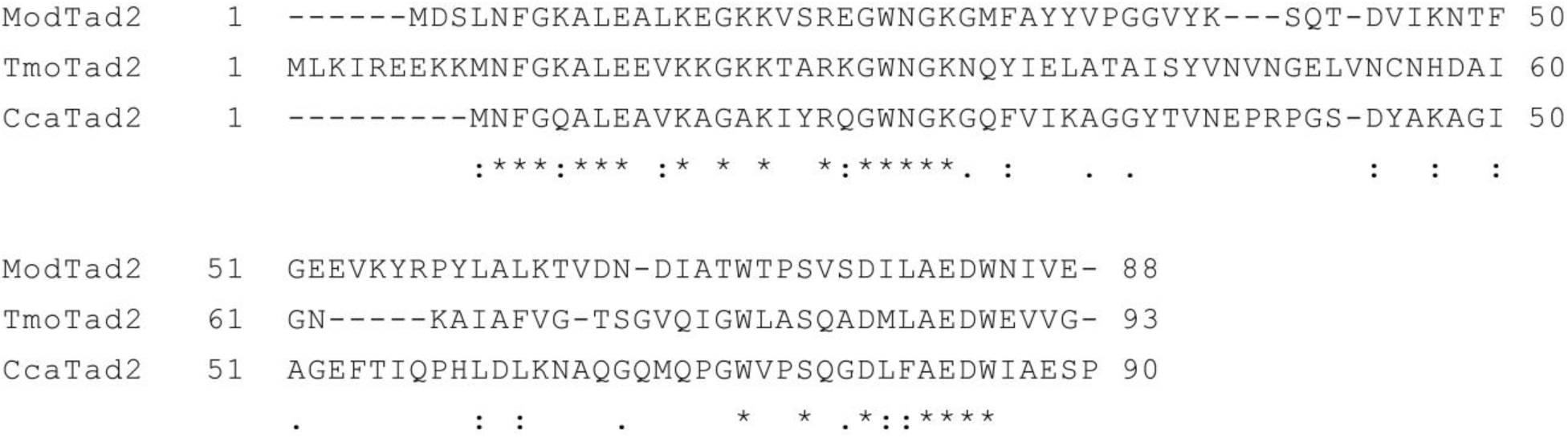
Multiple sequence alignment of ModTad2, TmoTad2 and CcaTad2. Multiple sequence alignment was performed using the hhpred server^59^.

**Figure S3.**
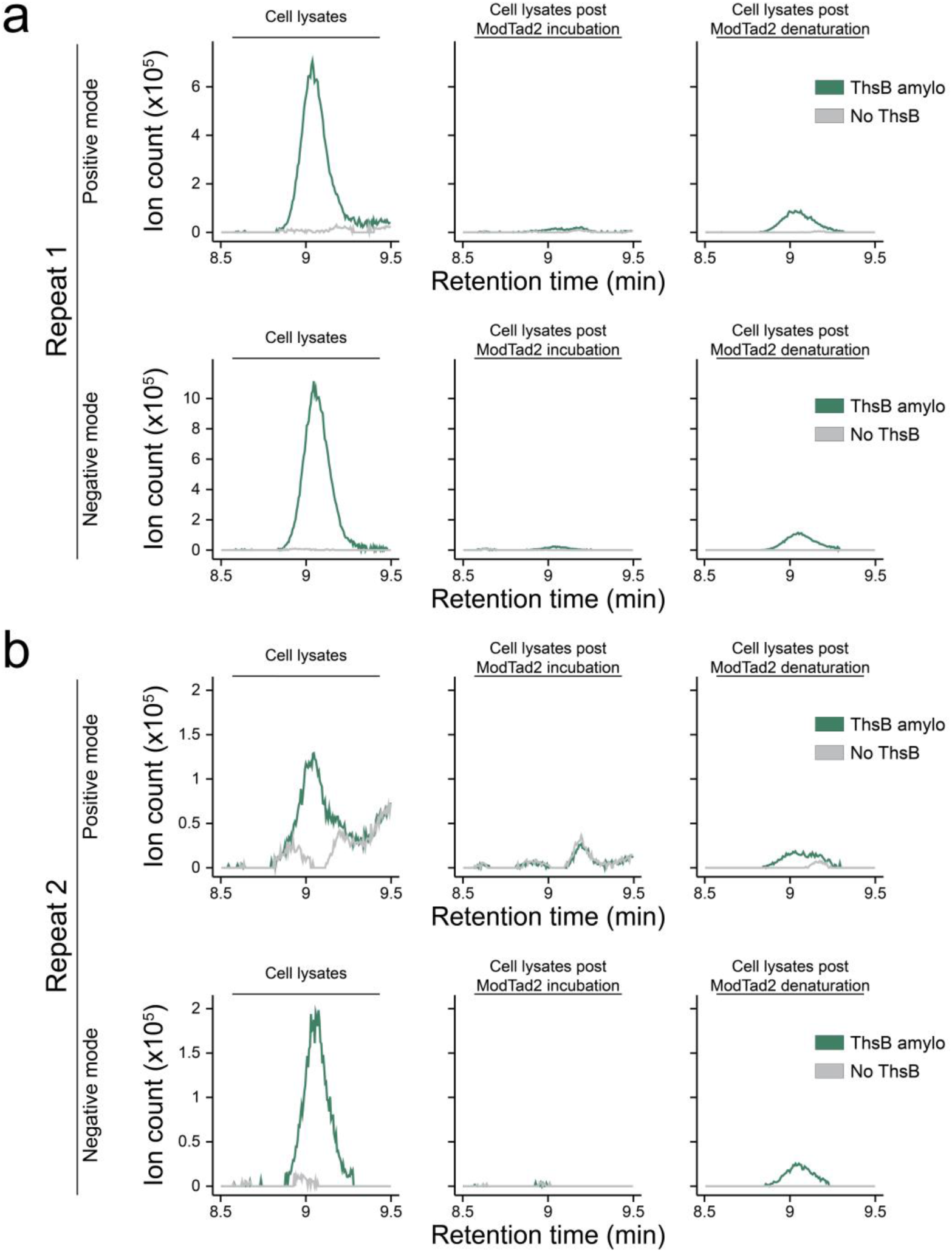
Purified ModTad2 protein binds type II Thoeris-derived signaling molecule. **a**, **b**, Two repeats of the mass spectrometry (MS) analysis shown in Figure 3a, analyzed in both positive and negative modes. Cells expressing ThsB_amylo_ or control cells that express GFP instead were infected with phage SBSphiJ at MOI = 10. After 120 min the cells were lysed and lysates filtered, then incubated with purified ModTad2. The lysates prior and post incubation with ModTad2, or after denaturation of ModTad2, were analyzed by mass spectrometry (MS). Masses visualized in positive mode are in the m/z range of 697.1262 – 697.1462 with a retention time of 8.5 – 9.5 min. Masses visualized in negative mode in S3 are in the m/z range of 695.1153 – 695.1353 with a retention time of 8.5 – 9.5 min. Extracted mass chromatograms of ions with an m/z value of 697.1374 and retention time of 9.04 min are presented in positive mode. In negative mode, extracted mass chromatograms of ions with an m/z value of 695.1251 are presented. Data in panel a are the same data presented in Figure 3a.

**Figure S4.**
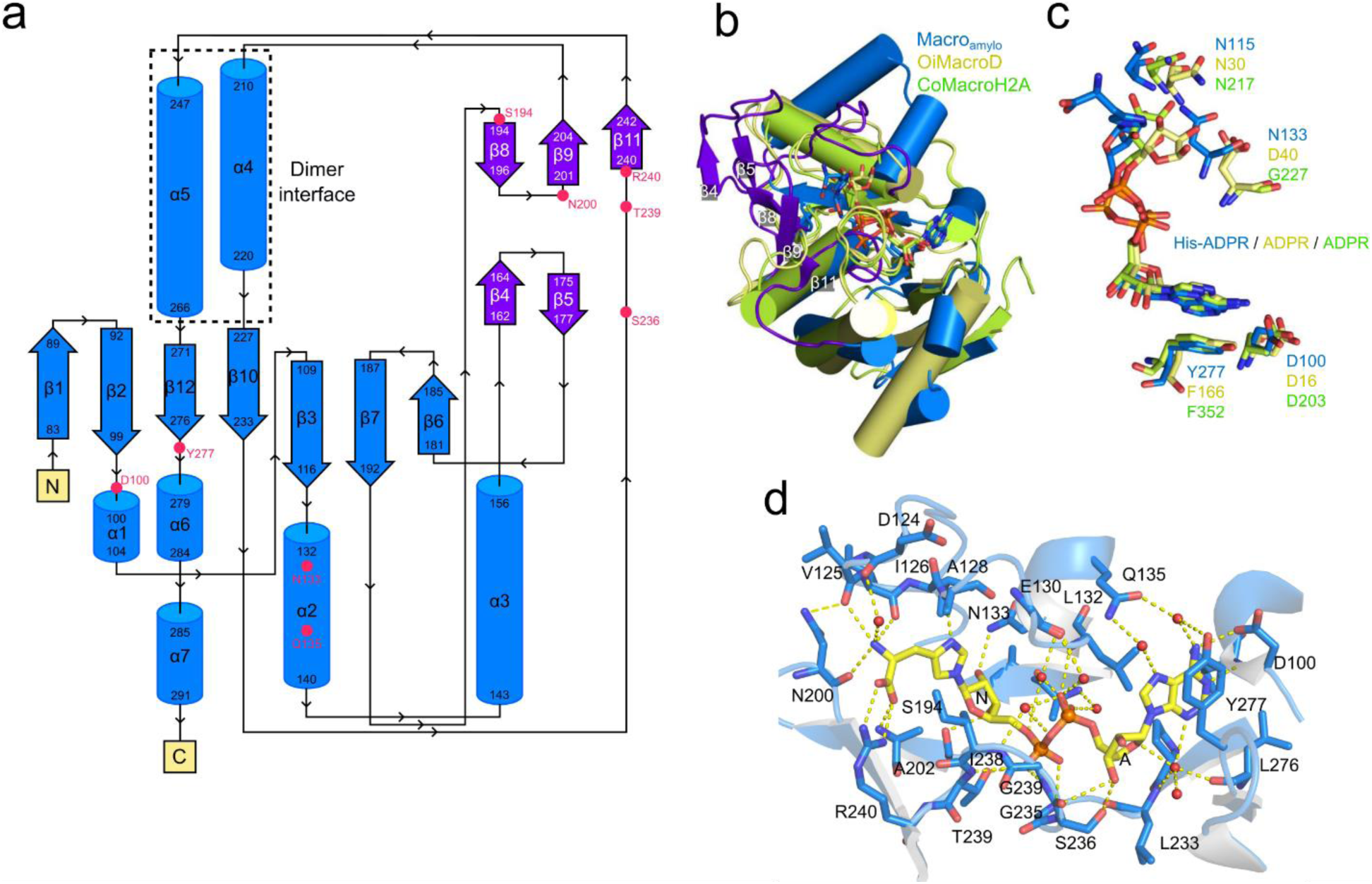
Structural features of the Macro_amylo_ domain. **a,** Topology diagram of Macro_amylo_ domain structure. Unique ThsA elements: a β hairpin (strands β4 and β5) and a small beta sheet (strands β8, 9, 11) are colored purple. Residues that make side-chain contacts to the ligand are marked by red circles. **b**, Superposition of the Macro_amylo_ domain (blue) with a ADPR-bound type OiMacroD domain of bacterium *Oceanobacillus iheyensis* (yellow, PDB ID 5LAU, Dali Z-score 15.3, r.m.s.d. 2.8 Å2 over 168 aligned atoms, 17% identity)^18^ and MacroH2A-like macrodomain from *Capsaspora owczarzaki* CoMacroH2A (green, PDB ID 7NY7, Dali Z-score 13.7, r.m.s.d. 2.7 Å^2^ over 152 aligned atoms, 13% identity)^19^. The Macro_amylo_ domain possesses longer loops near the binding pocket (colored purple). Unique ThsA elements: a β hairpin (strands β4 and β5) and a small beta sheet (strands β8, 9, 11) are marked. **c**, Conserved residues of the ligand binding pockets of the Macro_amylo_ (blue), OiMacroD (yellow) and CoMacroH2A (green). Ligand molecules bound in the binding pocket are colored respectively. MacroD catalytic aspartate D40 is not conserved in ThsA and CoMacroH2A. **d**, Detailed view of the ThsA residues interacting with His-ADPR. Yellow dashed lines denote hydrogen bonding interactions. A and N marks A- and N-ribose, respectively.

**Table S1. MSMS data or His-ADPR molecule released from denatured ModTad2, in positive ionization mode (provided as an Excel sheet)**

**Table S2. Strains of bacteria and phages used in this study (provided as an Excel sheet)**

**Table S3. Plasmids and primers used in this study (provided as an Excel sheet)**

**Table S4.** Summary of data collection and refinement statistics.

|  | Macro<sub>amylo</sub>*<br>His-ADPR bound<br>(8R66) | ModTad2<br>apo*<br>(8V3E) |
| --- | --- | --- |
| **Data collection** |  |  |
| Space group | C 1 2 1 | H 3 2 |
| Cell dimensions<sup>21</sup> |  |  |
| *a*, *b*, *c* (Å) | 140.30, 71.06, 51.96 | 93.21, 93.21, 235.55 |
| $\alpha$, $\beta$, $\gamma$ (°) | 90, 96.78, 90 | 90.00, 90.00, 120.00 |
| Resolution (Å) | 69.7–2.23 (2.34–2.23) | 47.57–2.39 (2.48–2.39) |
| *R*<sub>merge</sub> | 0.057 (0.366) | 0.094 (0.177) |
| *R*<sub>pim</sub> | 0.025 (0.198) | 0.039 (0.071) |
| *I* / σ*I* | 9.0 (1.8) | 11.9 (6.5) |
| Completeness (%) | 97.1 (79.9) | 99.0 (95.2) |
| Redundancy | 6.5 (4.4) | 6.9 (6.6) |
| **Refinement** |  |  |
| Resolution (Å) | 69.7–2.23 | 45.57–2.39 |
| No. reflections |  |  |
| Total | 159305 | 109160 |
| Unique | 24248 | 15786 |
| Free | 2454 | 1507 |
| *R*<sub>work</sub> / *R*<sub>free</sub> | 0.186 / 0.226 | 0.227 / 0.273 |
| No. atoms |  |  |
| Protein | 3435 (2 copies) | 2503 (4 copies) |
| Ligand/ion | 97 | – |
| Water | 82 | 206 |
| *B*-factors |  |  |
| Protein | 55.62 | 25.61 |
| Ligand/ion | 49.93 | – |
| Water | 52.44 | 26.13 |
| R.m.s. deviations |  |  |
| Bond lengths (Å) | 0.003 | 0.002 |
| Bond angles (°) | 0.598 | 0.440 |

\*Dataset was collected from an individual crystal Values in parentheses are for the highest-resolution shell.

